# Migrasomes May Originate from Cilia and Represent an Alternative Ciliary Form

**DOI:** 10.64898/2026.10.06.757022

**Authors:** Siyu Zhang, Abudurexiti Kutibiding, Mengci Wang, Yi Yang

## Abstract

**Background:** Migrasomes, the newly discovered vesicular organelles, have long been speculated to originate from membrane dynamics during cell migration. Here we present integrated imaging and multi-omics evidence supporting a ciliary origin of migrasomes. The shared vesicular trafficking pathways between ciliogenesis and migrasome biogenesis (trans-Golgi/endosomal systems) prompted us to investigate their spatiotemporal coupling mechanism in disease models.

**Methods:** Using high-resolution SEM and fluorescence confocal microscopy (FCM), we visualized cilium-like structures with enlarged tips adjacent to migrasome-like vesicles across multiple cell types. TMT-based quantitative proteomics was used to identify proteins shared among migrasomes from different cell types and examine their associations with ciliary proteins. DMBA-induced breast cancer and femoral fracture models were used to investigate the cilia–migrasome association during tumor metastasis and fracture healing.

**Results:** Bioinformatics analysis showed a highly positive incidence correlation between cilia and migrasome genes in 31 types of 7858 human normal tissues and organs and various cell types. In 32 cancer tissues, cilia and migrasome genes were still strongly but abnormally correlated. Signaling enrichment by ARL13B and TSPAN4 showed that cilia–migrasome association is related to epithelial-mesenchymal transition (EMT). By analyzing the ARL13B and PIGK expression, we found the abnormal cilia–migrasome association in metastatic breast cancer and positive cilia–migrasome association in fracture healing bone.

**Conclusion:** Our findings support a ciliary origin of migrasomes, pointing to a previously unrecognized pathway of organelle biogenesis. Notably, the ciliary-migrasomal association correlates with two distinct pathological/physiological contexts: (1) aberrant epithelial-mesenchymal transition (EMT) in metastatic breast cancer, and (2) accelerated regeneration during fracture healing and tissue repair. This dual correlation not only establishes a mechanistic link between ciliary function and migrasome biogenesis but also highlights their divergent roles in disease progression versus tissue homeostasis. Investigating the spatiotemporal dynamics of this association may provide novel therapeutic targets for metastatic malignancies and regenerative medicine.

**Highlights:**

1. The Ciliary Origin of Migrasomes: Redefining Organelle Biogenesis in Development and Disease.
2. The Cilia and Migrasomes Structural and Functional Parallels
3. Rethinking Migrasome Biogenesis from a Ciliary Perspective.

## 1 Introduction

Migrasomes – the enigmatic vesicular organelles discovered in migrating cells – have long been shrouded in mystery regarding their cellular origin[1]. Migrasomes have been shown to regulate organ formation, cell migration, and mitochondrial quality control [2, 3], and play an important role in development and disease progression [4–6].While previous studies predominantly attributed migrasome biogenesis to retraction fiber dynamics during cell migration, emerging evidence reveals striking parallels between ciliogenesis and migrasome formation, particularly in their shared dependence on trans-Golgi and recycling endosomal vesicles[7]. This convergence of vesicular trafficking pathways compels us to re-examine the fundamental question: Could cilia be the progenitor of migrasomes?

Cilia, the microtubule-based signaling hubs, orchestrate developmental patterning through conserved pathways like Hedgehog and GPCR signaling [8, 9]. Their assembly requires precise coordination of centriole-to-basal body transition and vesicular docking – processes remarkably mirrored in migrasome formation[10–13]. Notably, both organelles:1. Rely on Rab GTPase-mediated vesicular transport (Rab8a/Rab11a in migrasomes vs. Rab35 in cilia) [7]; 2. Utilize SNARE complexes for membrane fusion (VAMP2/3 in migrasomes vs. SNAP25 in cilia)[7]; 3. Exhibit polarized secretion patterns (migrasome-mediated directional signaling vs. ciliary morphogen gradients)

The prevailing TSPAN4-centric model posits that migrasome formation is driven by tetraspanin-enriched macrodomains. However, our bioinformatics profiling reveals migrasome marker expression correlates with cilia-related gene set scores.This challenges the notion of migrasomes as mere byproducts of cell migration, instead suggesting a ciliary origin through vesicular remodeling.

Abnormal formation and function of cilia cause diseases related to signal transduction disorders (called “ciliopathy”), affecting most tissues or organs, including bone, brain, heart, kidney and liver, and also participate in the progression of cancer [14, 15].Ciliopathies spanning bone malformations to tumor progression exhibit striking overlaps with migrasome-related pathologies. Our preliminary data demonstrate dynamic cilia-migrasome coupling during fracture repair and pathological decoupling in metastatic breast cancer. These findings link the cilia–migrasome association to EMT and highlight its relevance to tumor progression and tissue repair.

In this study, for the first time, we investigated the association between migrasome and cilia genes by bioinformatics analysis and confirmed their association and the possibility of migrasomes release from cilia by cellular electron microscopy imaging and protein composition and correlation analysis. In addition, we showed for the first time that the cilia–migrasome association changes during fracture healing and tumorigenesis, which may provide new ideas for exploring the role of cilia–migrasome association in tissue regeneration and repair.

## 2 Materials and Methods

### 2.1 Reagents and antibodies

The used reagents included DAPI (CM07245, proteintech, USA), Ciliobrevin D (M14923, AbMole, Shanghai, China), and Dynarrestin (M14924, abmole, AbMole, Shanghai, China). The antibodies included ARL13B (17711-1-AP, proteintech, USA), Acetyl-Tubulin (66200-1-Ig, proteintech, USA), TSPAN4 (bs-9413R, Bioss, China), PIGK (sc-398611, Santa Cruz Biotechnology, USA), and CPQ (16601-1-AP, proteintech, USA).

### 2.2 Bioinformatics analysis

#### Spearman correlation analysis of cilia-related gene set scores and migrasome-related markers

From the UCSC Xena dataset (https://xenabrowser.net/datapages/), we downloaded gene expression matrixs of normal tissue samples and tumor tissue samples from the GTEx dataset and TCGA dataset, respectively. In R software, we constructed a Seurat object related to GTEx gene expression using the Seurat package and clustered them. Using the singscore algorithm of the irGSEA package and the gene sets of seven main cilia-related functions from the Gene Ontology dataset (GO, https://www.geneontology.org/), namely “Intraciliary Transport Involved In Cilium Assembly (GO:0035735)”, “Non-Motile Cilium Assembly (GO:1905515)”, “Motile Cilium Assembly (GO:0044458)”, “Cilium Disassembly (GO:0061523)”, “Cilium Movement (GO:0003341)”, “Cilium Organization (GO:0044782)”, and “Cilium-Dependent Cell Motility (GO:0060285)”, we scored the cilia function for each sample. Spearman correlation analysis was performed between the expression of migrasome-related markers (“NDST1”, “EOGT”, “PIGK”, “CPQ”, “TSPAN4”, “TSPAN7”) and the main cilia-related gene set score, respectively.

The single-cell RNAseq datasets of human normal tissues, including epithelial cells, endothelial cells, immune cells, and stromal cells, were downloaded from Tabula Sapiens (https://tabula-sapiens.sf.czbiohub.org/). In R software, the h5ad objects in the dataset were converted into Seurat objects by using Seurat and anndataR software packages, respectively, and then received batch effect correction and re-clustering using the harmony software package. Spearman correlation analysis was performed between the expression of migrasome-related markers (“NDST1”, ‘EOGT”, “PIGK”, “CPQ”, “TSPAN4”, “TSPAN7”) and the main cilia-related gene set scores, respectively.

#### Dynamic binding and dissociation analysis of ARL13B-TSPAN4

The ARL13B-TSPAN4 docking experiment was conducted using HDOCK. The binding free energy was calculated by MM/GBSA on the HawkDock server. The amino acid residues of the protein were completed using SPDBV 4.10 software, and the complete structure of the protein was exported and saved in pdb format. In Gromacs 2024.4 software, molecular simulation of protein interaction was carried out under the AMBER99SB-ILDN force field. A water box was set up and the TIP3P water model was used to simulate the addition of NaCl ions to saline for charge balance. Then, the steepest descent method was used for energy minimization. The system was equilibrated for 1 ns at 300 K to obtain a stable system. After equilibration, a 200-ns molecular dynamics simulation was performed. The trajectory file after simulation was subjected to translational elimination, and the simulation system was analyzed. The indicators of RMSD, RMSF, RG, SASA, and DSSP were analyzed to evaluate the effect of protein interaction.

### 2.3 Cell culture

Mesenchymal stem cells (MSCs) (No. CP-M131; Pricella, Wuhan, China), SH-SY5Y cells (iCell-h187), human skin fibroblasts (HSF) cells (HUM-iCell-s001; Icellbioscience, Shanghai, China) were cultured in MEM supplemented with 10% FBS. RAW264.7 cells (ICELL-M047, Icellbioscience, Shanghai, China), and HUVEC cells (iCell-h110; Icellbioscience, Shanghai, China) were cultured in DMEM (Thermofisher, USA) supplemented with 10% FBS. All cells were cultured in a cell incubator at 37 °C and 5% CO_2_.

### 2.4 Migrasome purification

Cells that were seeded on fibronectin-coated dishes were grown overnight and then the cells were washed with phosphate buffer solution (PBS). After digestion with trypsin, cells were centrifuged at 1000×g for 10 minutes to discard cell bodies, followed by centrifugation at 4000×g for 20 minutes to remove cell debris and centrifugation at 18000×g for 40 minutes to harvest crude migrasomes precipitate. The precipitate was resuspended in 800 μl sample buffer (400 μl extraction buffer + 400 μl 10% OptiPrep) and grading at 150000×g for 4 hours in an OptiPrep density gradient using an MLS50 rotor (Optima MAX-XP). The gradients were as follows: 5%, 10%, 15%, 20%, 25%, 30%, 35%, 40%, and 50%. Preparation fractions were used for negative staining, western blot analysis, BCA assay, or RNA extraction.

### 2.5 Scanning electron microscope (SEM)

Cells were fixed with 1ml of 2.5% glutaraldehyde at room temperature for 2 hours and then were rinsed 3 times with 0. 1M phosphate buffer PB (PH7.4) for 15 minutes each time, followed by fixtion with 1% osmic acid that was prepared with 0.1M phosphate buffer PB (PH7.4) in the dark at room temperature for 1-2 hours. The fixted cells were rinsed 3 times with 0.1M phosphate buffer PB (PH7.4) and subjected to dehydration with gradient alcohol (30%-50%-70%-80%-90%-95%-100%-100%, 15 minutes in each gradient) and isoamyl acetate (15 minutes). Cells were dried in a critical point dryer. The cell sample that was closely attached to the conductive carbon film double-side glue was sprayed gold for 30 seconds for conductive treatment on the sample stage of the ion sputtering instrument. The images were observed under a scanning electron microscope.

### 2.6 L929 cells stably expressing TSPAN4-GFP

L929 cells stably expressing TSPAN4–EGFP were a gift from Li Yu’s laboratory at Tsinghua University and generated through L929 cells transducing with the PiggyBac Transposon Vector System constructing with TSPAN4-EGFP (pB-TSPAN4-EGFP transposon vector) and pBASE (transposase vector) at a ratio of 1:3. After 24 hours of transduction, the cells were cultured in medium containing 600 mg/ml G418 or 200 mg/ml hygromycin B for 3-5 days. Single-cell sorting was performed using a fluorescence-activated cell sorting. The single-cell clone was observed using confocal microscopy.

### 2.7 Confocal microscopy

Cells were cultured in a confocal dish that was pre-coated with fibronectin (10□µg□ml-1) with or without drug treatment. After 16 hours of culturing in a 5% CO_2_ incubator at 37□, confocal dishes were placed on the Nikon A1 confocal microscope (Japan) to photograph the images.

### 2.8 Immunofluorescent staining

The HSF, HUVEC, and MSCs in the logarithmic phase were washed by PBS. Cell fixation was achieved by 5 minutes of immersion in the medium supplemented with 50% of 4% paraformaldehyde and 5 minutes of immersion in 4% paraformaldehyde. The fixed cells were subjected to 30 minutes of permeabilization and blocking treatments using 0.05% saponin and 10% FBS. Primary antibodies against ARL13B or acetylated Tubulin were diluted in PBS containing 10% FBS and used to incubate cells at 4°C overnight. After washing with PBS three times, cells were stained with the secondary antibody (diluted by PBS containing 10% FBS) for 1 hour. The cells were washed with PBS three times and then counterstained by WGA. Immunofluorescence was observed under confocal microscopy (A1/SIM/STORM, Nikon).

For tissue immunofluorescence, the paraffin sections were deparaffinized and rehydrated, then placed in antigen repair buffer (G1202, PH 6.0, citric acid) for antigen repair. Tissue sections were drip-added with 3% hydrogen peroxide solution and incubated at room temperature in the dark for about 25 minutes. After being washed three times with PBS, the sections were immunofluorescence reacted with anti-ARL13B (proteintech 17711-1-AP), anti-PIGK (SATNTA sc-398611), and CPQ (proteintech 16601-1-AP) in sequence, in a repeated procedure that included serum blocking for 30 minutes, the primary antibody incubation overnight, HRP-labeled secondary antibody incubation, fluorescent labeling (iF488-TSA staining, iF555-TSA, iF647-TSA, respectively) and antibody elution (G1266). The nuclei were counterstained with DAPI. The sections were sealed with an anti-fluorescence quench sealing agent.

For migrasome immunofluorescence, migrasomes were fixed with 4% paraformaldehyde for 15 to 30 minutes. The migrasomes were permeabilized with 0.1% Triton X-100 for 5 min and then blocked by 5% goat serum for 30 minutes, followed by primary antibody incubation overnight at 4 °C. After washed once with TBST for 5 minutes and twice with TBS for 5 minutes each, migrasomes were incubated with fluorescent secondary antibody for 1 hour at 37 °C in the dark. After washing once with TBST for 5 minutes and twice with TBS for 5 minutes each, migrasomes were counterstained with WGA dye. After adding the anti-fluorescence attenuation sealant, the migrasome images were observed under a fluorescence microscope.

### 2.9 Sample preparation and quantitative mass spectrometry

Protein sample was prepared through lysing migrasomes in lysis buffer (diluted in PBS) containing 8M urea and protease inhibitors (BioTools, Jupiter, Florida). Protein samples with equal amounts were reduced, alkylated, and digested with trypsin and labeled with TMT reagent (Thermo, Pierce Biotechnology, USA). The generated TMT-labeled peptides were desalted with a Sep-Pak C18 column. Tryptic peptides were divided into 12 fractions using a high-pH RPLC (XBridgeTM BEH300 C18 5μm, 300A, inner diameter 250 mm × 4.6 mm, Waters).

For LC-MS/MS analysis, TMT-labeled peptides were separated via A C18 column (150 mm length, 75 µm inner diameter, 5 µm, 300 A) with a Thermo-Dionex Ultimate 3000 HPLC system, a system was connected to a Fusion Lumos Tribrid mass spectrometer. A series of adjusted linear gradients were applied at a flow rate of 300 nL/min depending on the hydrophobicity of the fractions. A data-dependent acquisition mode was set in the mass spectrometer to acquire data. Survey scans ranged from m/z 375 to 1550. After a single microscan, the first N strongest peaks with charge states 2 and above were separated by a normalized collision energy of 35%. The MS2 spectra were with a maximum injection time of 60ms, an AGC target of 1e5, and a resolution of 17,500. The isolation window was set to 0.7 Da width and the dynamic exclusion time was 60s..

MS/MS spectra were generated through searching Uniprot Rat dataset (January 10, 2015; 89105 sequence) by the SEQUEST search engine in the Proteome Discoverer 1.4 software. The search criteria were as follows: Full trypsin specificity, allowed one missed cut, urea methylation on cysteine and TMT hexaomer on the N-terminus of lysine/peptide being set as fixed modifications, precursor ion mass tolerance being set to 10 ppm for all MS, oxidation on methionine being set as variable modifications, and the fragment ion mass tolerance being set to 20 mmu for all MS2 spectra. Peptide profile matches (PSM) were verified with a 1% false discovery rate (FDR) based on q values using the Percolator provided by Proteome Discoverer software. Proteomic analyses were performed in triplicate. A ratio >0.75 or <1.3 was considered to be a differentially expressed protein when the p-value was < 0.05. Proteome data were deposited into the ProteomeXchange Consortium through the PRIDE Partner Repository under the dataset identifier PXD020522.

### 2.10 Mice and rats

All animal experiments were approved by the Animal Ethics Committee Laboratory of the First Affiliated Hospital of Xinjiang Medical University and performed according to the guidelines of the National Health and Medical Research Council of China. C57/BL6 mice (male, 8-week-old) with an average body weight of 20 g and female Sprague Dawley (SD) rats (4-5-week-old, weight of 85-130g) were obtained from Xinjiang Medical University and housed in individual cages with a 12 h light/dark cycle and fed standard food and water at 22-28C and 60%-70% relative humidity.

### 2.11 Mouse model of fracture

Mice were anesthetized through an intraperitoneal injection of 1% pentobarbital. Animals were fixed in the lateral decubution position with skin preparation and routine disinfection of the outer thigh. The skin was cut 1cm along the femur to expose the vastus medialis muscle and knee joint. The muscle was bluntly separated in the middle of the femur and the femur was free for 5mm. The femur was sawed off with a dental drill configured with a disc bit in the middle of the long axis of the femur. A spherical drill was used to drill through the bone marrow cavity from the distal end of the femur along the long axis of the femur, and then the bone marrow cavity was pulled out. The pulpal opening was cooled down with precooled saline. The internal needle was inserted into the channel of the medullary cavity and drilled to a depth of approximately 3 mm. The sawed femur was connected and secured with an intramedullary nail. The surgical skin incision was sutured.

### 2.12 Establishment of a rat 7, 12-Dimethylbenz[a]anthracene (DMBA) model of breast carcinoma

Rats were given a single subcutaneous injection of DMBA solution (5 g DMBA dissolved in 500 mL sesame oil) at 100mq/kg body weight to induce breast tumors as demonstrated in previous studies [16, 17]. After injection, the rats were fed with standard food and water (12 h light/dark cycle) until 24 weeks old. Rat breast carcinoma in situ and metastatic tissues were obtained for further study.

### 2.13 Statistical methods

Statistical analysis was performed in GraphPad Prism (Version 7.0). All data were obtained from at least three independent experiments. The error line in the figure represents the standard deviation (SD). The t-test was used to compare the differences between groups.

## 3 Results

### 3.1 Cilia-migrasome has a strong and positive incidence relation in a variety of healthy tissues and cells

We investigated the incidence relation between cilia and migrasome via Spearman correlation analysis of cilia-related gene set scores and migrasome markers in normal tissues and found that NDST1, EOGT, PIGK, CPQ, TSPAN4, and TSPAN7 were highly positively correlated with cilia-related gene set scores (**Supplemental Data 1**). To exclude the strong cilia-migrasome correlation in a single tissue or organ in improving the overall cilia-migrasome correlation, we analyzed the correlations of cilia and migrasome in 31 tissue and organ separately (**Figure 1**). The results showed that except in the fallopian tube, testis, and uterus, which play main functions by transporting materials through ciliary movement, cilia, and migrasomes were highly negatively correlated, while in other organs, cilia and migrasomes were highly positively correlated (**Figure 1**) suggesting that there are significant organ differences in the incidence relation between cilia-migrasomes. However, in skin tissues, migrasome genes were positively correlated with non-motile ciliary assembly, ciliary transport involved in ciliary assembly, and ciliary organization but negatively correlated with motor ciliary assembly, ciliary motility, ciliary disassembly, and cilium-dependent cell motility (**Figure 1**) suggesting that differences in ciliary function in the same tissue also affect the association form of cilia and migrasomes. To further explore the cilia–migrasome association, we downloaded single-cell sequencing data about normal epithelial cells, endothelial cells, immune cells and stromal cells and analyzed the cilia–migrasome association in these cells. The results were consistent with the above analysis and found a highly positive correlation between cilia-related gene set scores and migrasome-related markers in epithelial cells (**Figures 2A-D**) as well as in endothelial cells, immune cells and stromal cells (**Supplemental Data 2-4A-D**). These data indicate a positive cilia–migrasome association in various normal cells.

**Figure 1.**
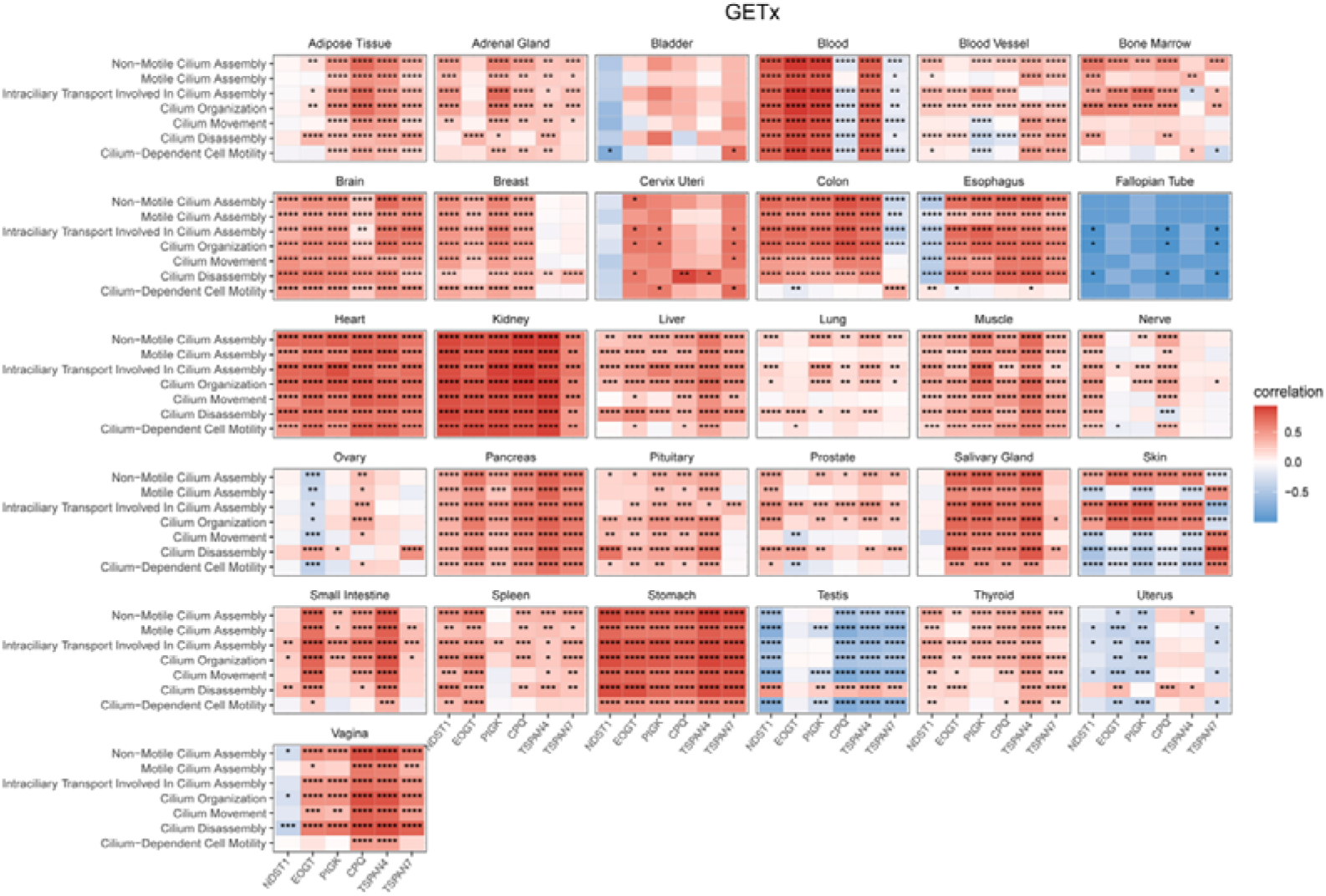
Cilia–migrasome association in various healthy tissues and cells. Spearman correlation analysis was performed between the expression of migrasome-associated markers (“NDST1”, “EOGT”, “PIGK”, “CPQ”, “TSPAN4”, and “TSPAN7”) and the ciliary major functional gene set scores, respectively, in single normal tissue of 31 different normal tissue samples.

**Figure 2.**
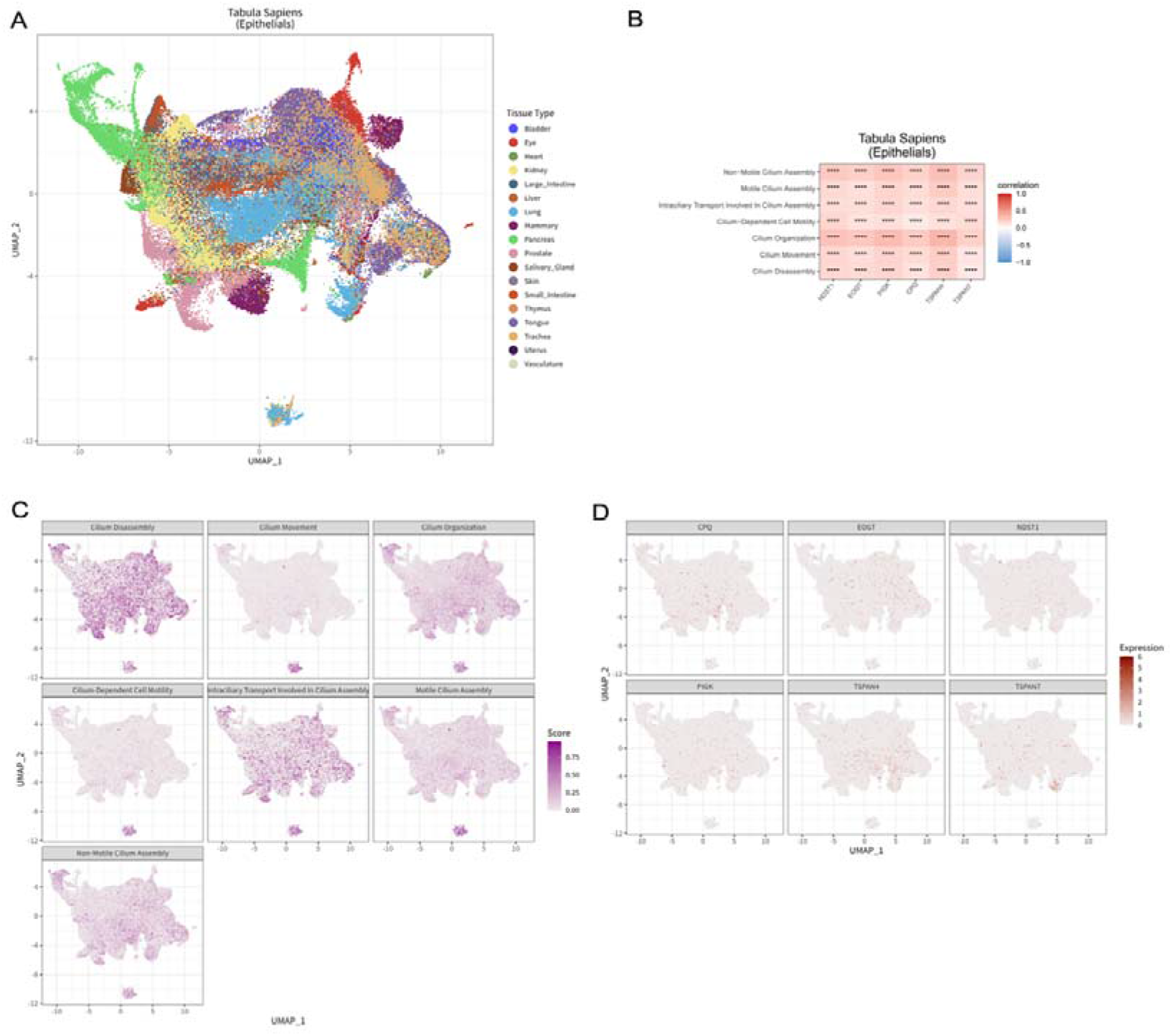
**Cilia–migrasome association in epithelial cells**. (A) UMAP plot of epithelial cells derived from normal tissues and organs. (B) Spearman correlation analysis was performed between the expression of migrasome-associated markers (“NDST1”, “EOGT”, “PIGK”, “CPQ”, “TSPAN4”, and “TSPAN7”) and the ciliary major functional gene set scores, respectively. (C) UMAP plot displaying cilium major functional gene set scores distributing in epithelial cells. (D) UMAP plot displaying migrasome-associated markers in epithelial cells.

### 3.2 The correlation between cilia and migrasome is abnormal in various tumor tissue

To further explore whether cilia–migrasome association in the pathological condition is consistent with that in normal tissues, we performed Spearman correlation analysis on cilia-related gene set scores and migrasome markers. The results showed that cilia-related gene set scores were positively correlated with TSPAN4 and negatively correlated with EOGT. NDST1 negatively correlated with non-motor ciliary assembly, motor ciliary assembly, and cilium-dependent cell movement. PIGK was negatively correlated with ciliary disassembly. CPQ was negatively correlated with cilium-dependent cell motility, and TSPAN7 was negatively correlated with ciliary transport involved in ciliary assembly (**Supplemental Data 5**). By analyzing the cilia–migrasome association in each tissue and organ separately, we found that the correlation between different migrasome markers and cilia-related gene set scores was quite different, which also indicated an abnormal incidence relation state of cilia and migrasome under pathological conditions (**Figure 3**). Our findings suggest that the abnormality of cilia–migrasome association may be one of the important mechanisms in tumorigenesis. However, whether these abnormal state changes affect the occurrence and development of tumors has not been studied.

**Figure 3.**
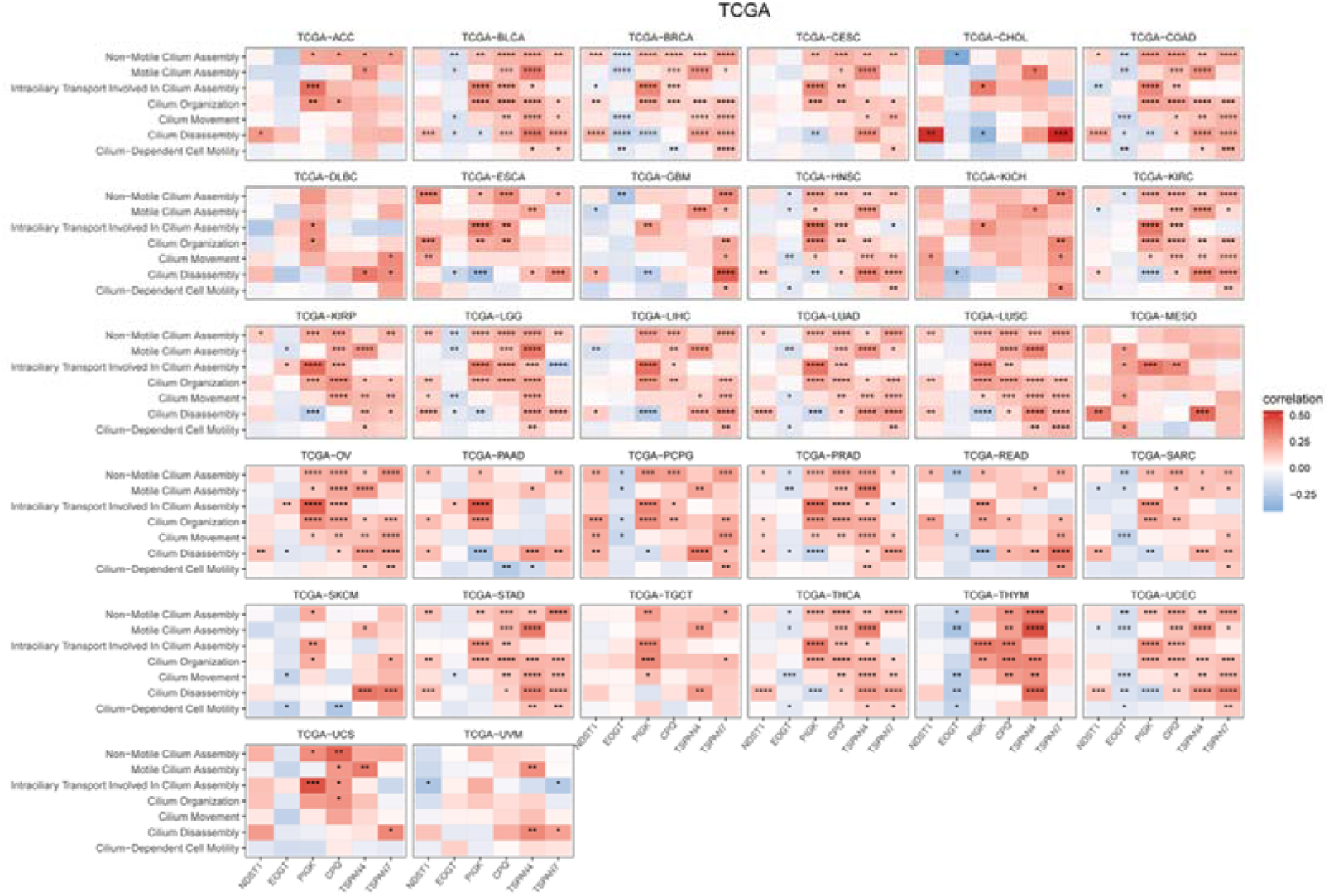
**Cilia–migrasome association in tumor tissues**. Spearman correlation analysis was performed between the expression of migrasome-associated markers (“NDST1”, “EOGT”, “PIGK”, “CPQ”, “TSPAN4”, and “TSPAN7”) and the ciliary major functional gene set scores, respectively in each kind of tumor of 32 kinds of tumor samples from TCGA dataset.

### 3.3 Cilia were enriched in the region where migrasomes were generated, and the cilium ends were enlarged

The morphology, distribution, and surface characteristics of the retraction fiber accompanying the migrasome and cilia were significantly different under SEM, making them easy to distinguish in SEM images. Specifically, the retraction fibers display dynamic and irregular protrusions structure, while cilia display regular and uniform structures (about 5-10 μm in length and 0.2μm in width). By observing the SEM images of RAW264.7 cells and SH-SY5Y cells, cilia-like structures distinguished from retraction fibers were enriched around the migrasome-shaped vesicular structures (about 2um), and the ends of the cilia-like structures enriched around the migrasome were enlarged, as if there were vesicular structures (**Figures 4A,D**). To further observe the association between cilia and migrasomes, we used SEM to observe the enrichment sites of migrasomes in various cells. The results showed that in RAW264.7 cells, mouse MSCs, SH-SY5Y cells, and HSF, cilia enriched in the production process of migrasomes and the end of the ciliary structure was enlarged around the migrasome, which seems to produce a vesicle-like structure similar to the migrasome (**Figures 4B,C,E,F,G-J**).

**Figure 4.**
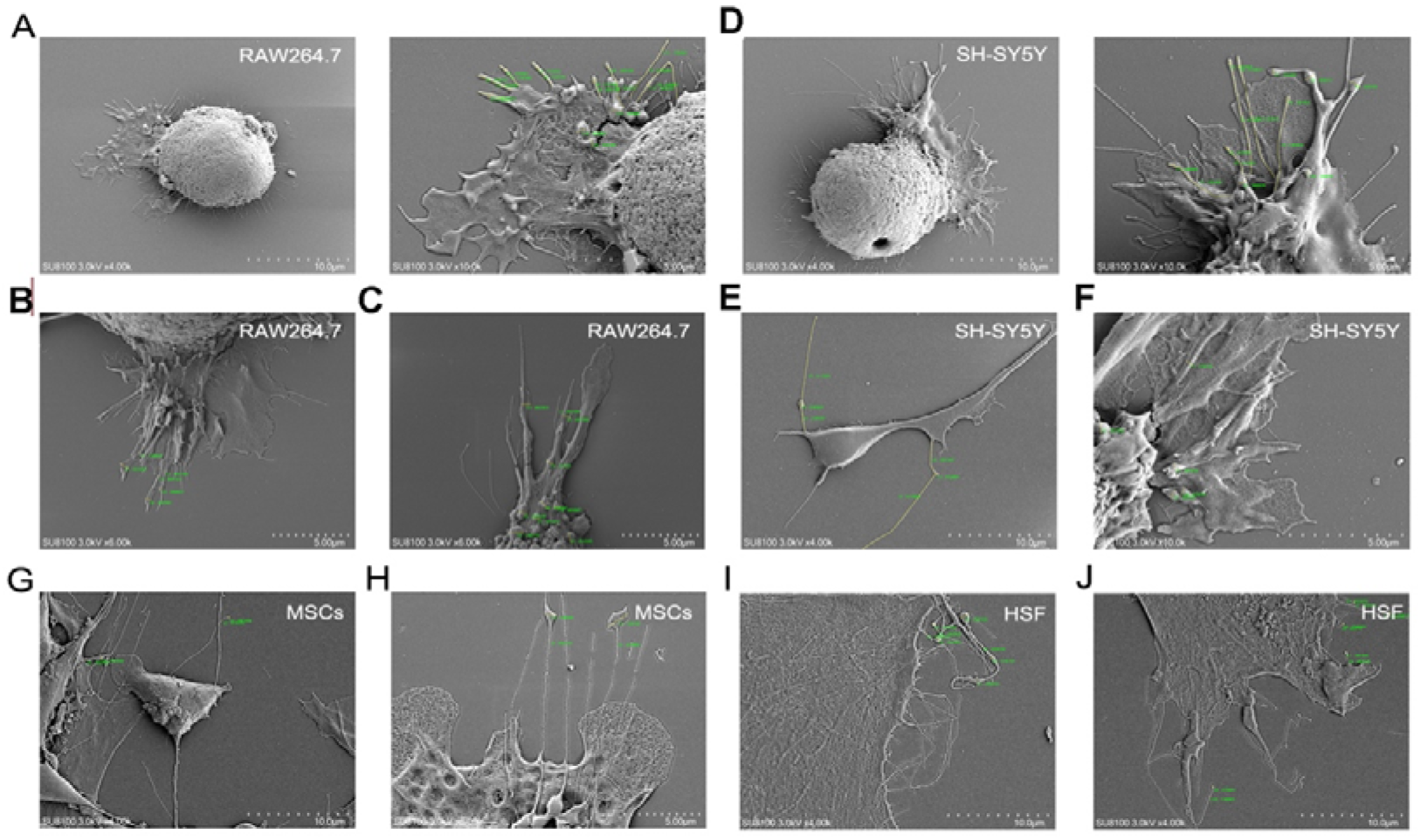
The relationship between migrasome production and cilia was observed by SEM in various kinds of cells. (A) SEM image of RAW264.7 cells. Scale bar, 10 μm. (B,C) SEM image displays migrasome at the tail of RAW264.7. Scale bar, 10 μm. (D) SEM image of SH-SY5Y cells. Scale bar, 10 μm. (E,F) SEM image displays migrasome at the tail of SH-SY5Y cells. Scale bar, 10 μm. (G) SEM image of MSCs. Scale bar, 10 μm. (H) SEM image displays migrasome at the tail of MSCs. Scale bar, 10 μm. (I) SEM image of HSFs. Scale bar, 5 μm. (J) SEM image displays migrasome at the tail of HSFs. Scale bar, 5 μm.

An externally sourced microscopy video illustrates dynamic remodeling of thin cell-associated protrusions bearing bead-like enlargements **(Supplementary Data 7; [37])**. These morphological dynamics motivate further investigation of transitions between rounded and elongated structures in the context of our model, which proposes that migrasomes may represent a vesicular form of cilia.

### 3.4 Ciliary marker proteins are detected in migrasomes

To further explore the cilia–migrasome association, we used immunofluorescence staining of ARL13B (a small GTPase that mainly serves as a cilium marker protein) or acetylated Tubulin (a critical protein for cilium formation and function and is highly enriched in cilia) on HSF, HUVEC, MSCs, and HSF, with WGA counterstain to highlight the overall structure, to observe whether ciliary marker proteins were expressed on the migrasomes. We found that cilium-marker proteins were expressed in migrasome in these cells, which also indicated the important cilia–migrasome association and effectively distinguished the relationship between retraction fibers and cilia in migrasome production (**Figures 5A-F**). To better demonstrate the expression of cilium marker proteins in the migrasome, we extracted the migrasome produced by cells and found that the migrasome still expressed ARL13B in the extracted migraosme (**Figures 5G, I**). This finding suggests that migrasomes express cilium marker proteins that not only appear as functional proteins during cell production but also remain expressed by isolated migrasome.

**Figure 5.**
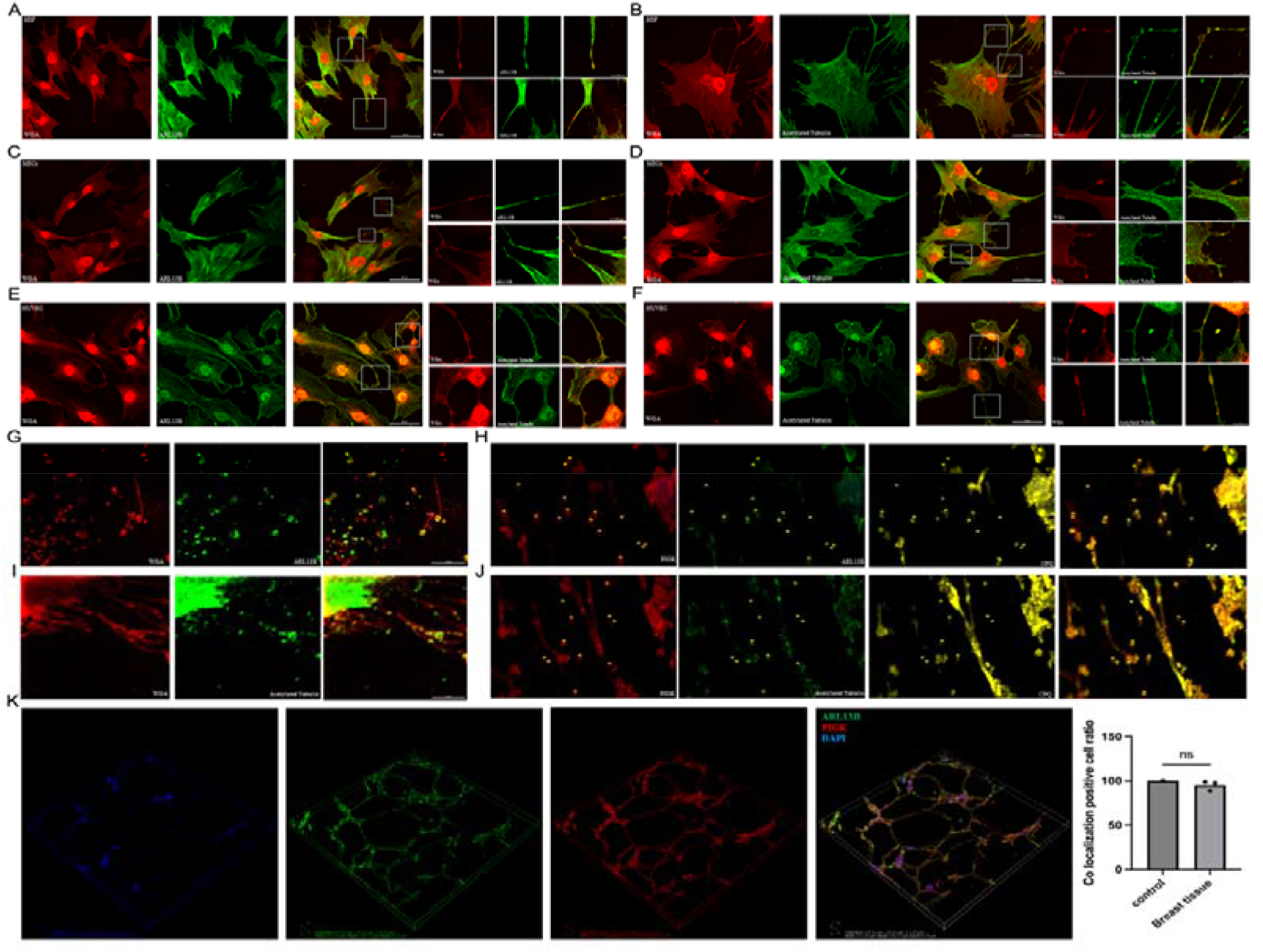
Immunofluorescence staining was used to observe the protein expression of migrasomes and cilia marker proteins in a variety of cells. (A,B) Expression of migrasome ARL13B (A) and Acetylated Tubulin (B) in HSF cells. Scale bar, 50μm. (C,D) Expression of migrasome ARL13B (C) and Acetylated Tubulin (D) in MSCs. Scale bar, 50μm. (E,F) Expression of migrasome ARL13B (E) and Acetylated Tubulin (F) in HUVEC. Scale bar, 50μm. (G) Expression of ARL13B in cell-extracted migrasomes. Scale bar, 10 μm. (H) PIGK, CPQ, and ARL13B were immunofluorescent stained in mouse mammary tissues and analyzed for vesicles ranging from 0.5 to 3um. Scale bar, 5 μm. (I) Expression of Acetylated Tubulin in cell-extracted migrasomes. Scale bar, 5 μm. (J) PIGK, CPQ, and Acetylated Tubulin were immunofluorescent stained in mouse mammary tissues and analyzed for vesicles ranging from 0.5 to 3um. Scale bar, 5 μm. (K) 3D confocal staining of PIGK and ARL13B in mouse mammary tissues.

To better observe the incidence relation between migrasome and cilia, evidence of their presence was examined in tumor tissues of breast cancer rats. We selected rat mammary tissues for immunostaining of migrasome markers (PIGK, CPQ) and ciliary marker (ARL13B), and analyzed vesicles from 0.5 to 3um. The results revealed that migrasome expressed ARL13B in the tissues (**Figures 5H, J**). To define the vesicular structure of migrasome and visualize the distribution of ARL13B in migrasome, we photographed them using confocal microscopy in the breast cancer tissues. Confocal imaging showed ARL13B co-distribution with PIGK-positive structures (Figure 5K). The phenomenon that migrasome still expressed the ciliary marker ARL13B in the complex microenvironment of tissues indicates that migrasome may be derived from cilia and that there is an important incidence relation between them.

### 3.5 Pharmacological and proteomic evidence for the cilia–migrasome association

Overexpression of TSPAN4 significantly enhanced the formation of migrasome and retraction fiber. To further explore the cilia–migrasome association, we examined the effects of Ciliobrevin D and Dynarrestin in L929 cells expressing TSPAN4–GFP. Treatment reduced TSPAN4-positive migrasome structures **(Figure 6A)** and TSPAN4 protein levels **(Figure 6B)**.

**Figure 6.**
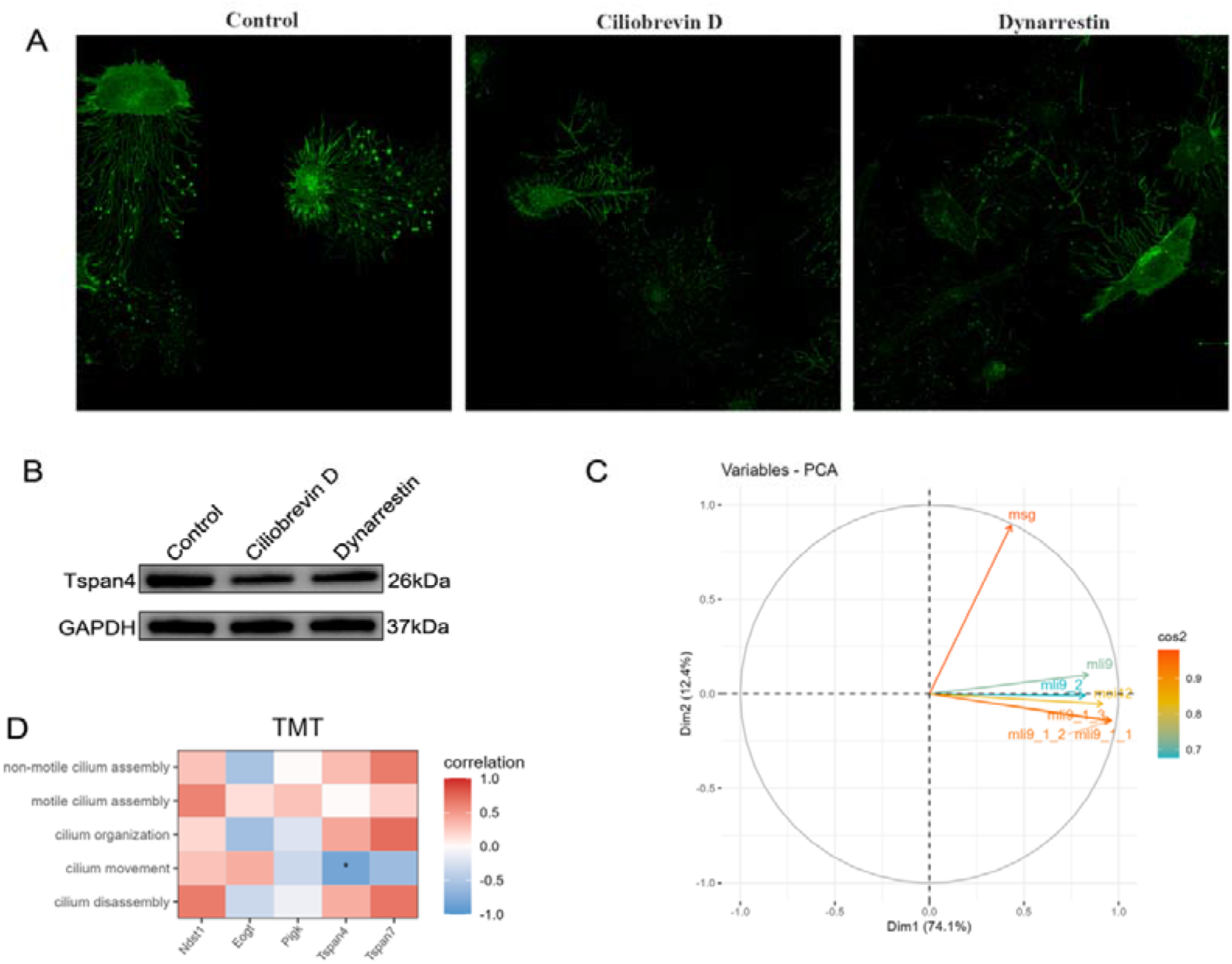
Pharmacological and proteomic evidence for the cilia–migrasome associations. (A) The migrasome production from TSPAN4-L929 cells cultured with Ciliobrevin D (30ug/ml) and Dynarrestin (30μg/ml) for 24h was observed by fluorescence confocal microscopy. Scale bar, 50μm. (B) The expression of TSPAN4 in L929 cells was detected by western blot. (C) PCA analysis of protein expression of migrasome TMT samples was performed using the FactoMineR and factoextra software packages. (D) The Spearman correlation analysis between migrasome markers (“NDST1”, “EOGT”, “PIGK”, “CPQ”, “TSPAN4”, and “TSPAN7”) and the ciliary major functional gene set scores.

To confirm the origin of the migrasome from cilia, we used TMT labeling to analyze the protein expression of migrasomes in multiple cells. Multiple samples were pooled for analysis to identify the common proteins expressed in multiple cell-derived migrasomes. PCA analysis of mouse samples was performed on protein expression of migrasome from TMT sample analysis using the FactoMineR and factoextra software packages. The results revealed similar protein expression in multiple cell-derived migrasomes (**Figure 6B**). However, we found strong differences between bone marrow mesenchymal stem cells-derived migrasomes and other cell-derived migrasomes, which led us to speculate that stemness might be related to migrasome composition. To explore the relationship between the components of migrasome proteins and cilia, we used the zscore algorithm of the GSVA software package, combined with the cilia-related functional gene set scores in the GO dataset described above, to score the ciliary function of the migrasome protein by TMT data, and analyzed the spearman correlation between migrasome markers and these functional scores (**Figure 6C**). The results were similar to those of the above studies, where migrasomes were strongly correlated with cilia and were most strongly correlated with ciliary motility (**Figure 6D**).

### 3.6 The cilia–migrasome association is linked to EMT

To explore the important signaling pathways and key molecules in regulating the cilia–migrasome association, we utilized the ssgsea algorithm of the GSVA software package and the hallmark gene sets from the MSigDB dataset (https://www.gsea-msigdb.org/gsea/msigdb) as the enrichment background to score hallmark pathways for both the GTEx and GDC-TCGA datasets. Spearman correlation analysis was conducted separately on the hallmark pathway scores and the expression levels of TSPAN4 and ARL13B. Pathways with significant correlations (with an absolute correlation value greater than 0.3 and a p-value less than 0.05) were selected for visualization. The results of obtaining the intersection of pathways with significant correlation between the two genes showed that there were 14 intersections in GTEx (**Figure 7A**) and 0 intersections in GDC-TCGA (**Figure 7B**). In GTEx, the significant pathways associated with ARL13B and TSPAN4 were enriched in Epithelial Mesenchymal Transition (EMT), Uv Response Dn, Tgf Beta Signaling, Angiogenesis, and Wnt Beta, Catenin Signaling, Notch Signaling, Apical Junction, and Hypoxia. Among them, EMT was the most relevant, and Tgf Beta Signaling, Wnt Beta, and Catenin Signaling were also confirmed to be involved in EMT (**Figure 7C**). Therefore, we hypothesized that EMT is an important signaling pathway regulating the cilia–migrasome association (**Figure 7C**). Notably, no hallmark pathways were significantly correlated with both ARL13B and TSPAN4 in the GDC-TCGA dataset under the specified criteria (Figures 7B, D). To further explore the relationship between the cilia–migrasome association and EMT, TSPAN4-GFP-tagged L929 cells were treated with EMT inhibitor-1, an EMT inhibitor. Using confocal microscope imaging system observation, we found that inhibitor-1 significantly inhibited the production of migrasome (**Figure 7E**). Moreover, we found that TSPAN4 expression was decreased after cell treatment with EMT inhibitor-1 (**Figure 7F**). These data suggest that cilia–migrasome association is related to EMT.

**Figure 7.**
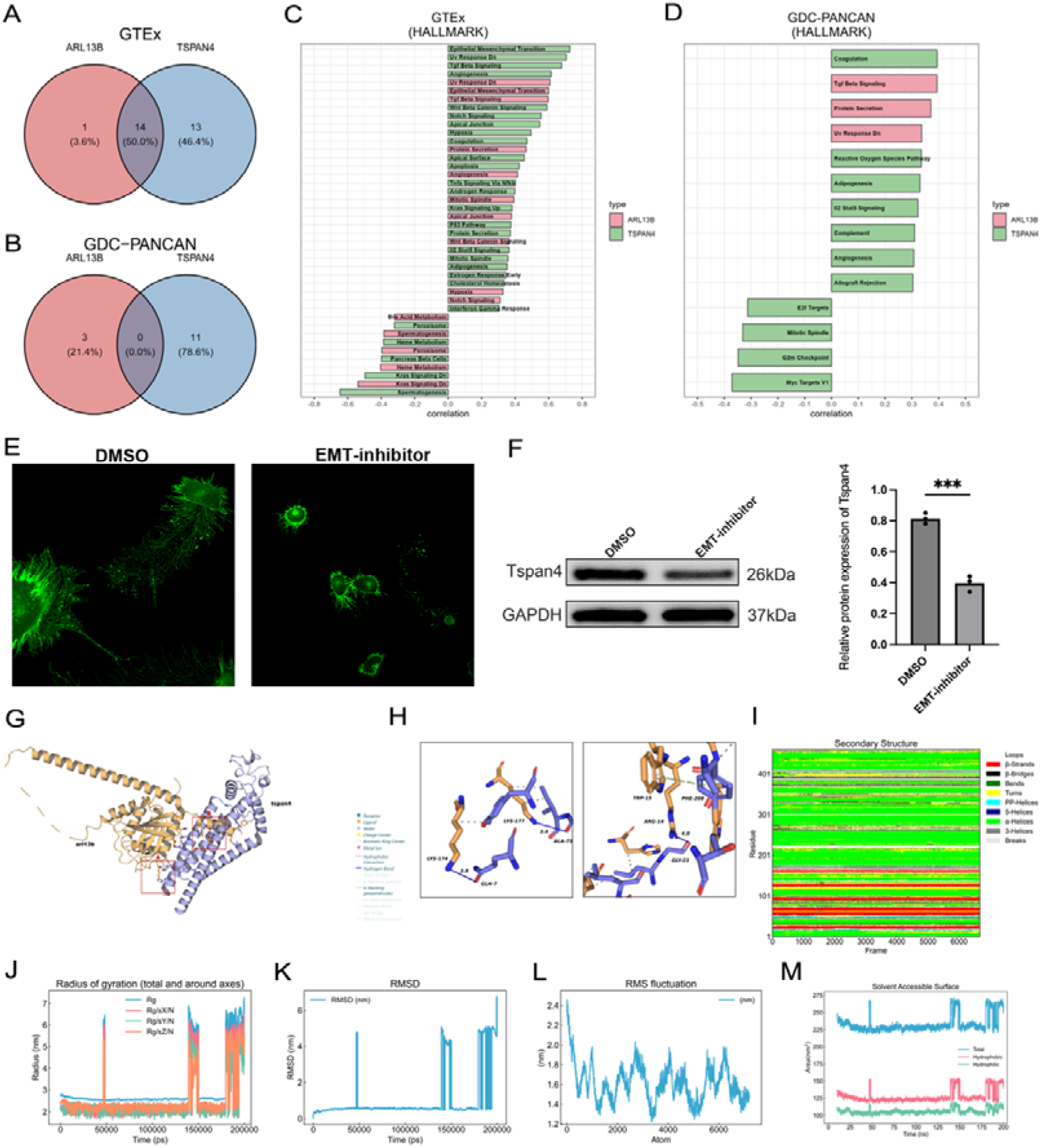
EMT as an important signal for the ciliary-migrasome incidence relation and the dynamic binding and dissociation between ARL13B-TSPAN4. (A,B) The significant associated pathway intersections between the two genes included 14 intersections in the GTEx dataset and 0 intersections in the GDC-TCGA dataset. (C,D) Visualization of significantly correlated pathways by the Spearman correlation analysis between the hallmark pathway score that was scored by ssgsea algorithm of GSVA software package and expression level of TSPAN4 and ARL13B (Absolute value of correlation greater than 0.3 and p value less than 0.05). (E) Laser confocal observation of migrator production in TSPAN4-L929 cells cultured with EMT-inhibitor (15ug/ml) for 24h. Scale bar, 50μm. (F) The expression of TSPAN4 in L929 cells treated with EMT-inhibitor (15ug/ml) for 24h was detected by western blot. (G) The interaction between ARL13B and TSPAN4 was analyzed by PLIP, and ARL13B was used as the reference strand for interaction analysis; yellow represents ARL13B and purple represents TSPAN4. Twenty-one pairs of hydrogen bonds are formed between the proteins. (H) A hydrogen bond is formed between LYS-174 of ARL13B and GLN-7 of TSPAN4 (solid blue line), and between LYS-177 of ARL13B and ALA-75 of TSPAN4. A hydrogen bond is formed between GLY-23 of TSPAN4 and ARG-14 of ARL13B, and a Pi-Pi cation interaction is formed between TRP-15 of ARL13B and PHE-208 of TSPAN4 (vertical, green). (I-M) The molecular dynamics (MD) simulation was performed for 200ns, and the simulated track files were translated and eliminated. Then the parameters of RMSD, RMSF, RG, SASA and DSSP were analyzed to evaluate the effect of protein interaction.

### 3.7 Computational modeling of ARL13B–TSPAN4 interactions

To further explore the incidence relation between cilia and migrasomes, the ARL13B-TSPAN4 docking experiment was conducted using HDOCK. The results showed that the lowest binding energy model_1 of ARL13B-TSPAN4 had a docking score of-396.45 and a confidence score of 0.9928 (**Figure 7G**), indicating a high possibility of ARL13B and TSPAN4 binding. The binding process of ARL13B and TSPAN4 was presented in the form of a surface (**Figure 7G**). The binding free energy calculated by the MM/GBSA method on the HawkDock server was-69.04 (kcal/mol) (**Figures 7G**, **H**). The results of molecular dynamics simulation analysis showed that 176 nm (88%) of the dynamic simulation docking presented a docking state, and 24 nm presented a non-docking discrete state (**Figures 7I-M**) indicating that the ARL13B-TSPAN4 is a dynamic binding and dynamic separation process with 88% of the time in a binding state and 12% of the time in a separated state. Since migrasomes are vesicular organelles that are shed and left behind in the original location during cell migration and released into the extracellular space, we speculate that the dynamic binding and separation process of ARL13B-TSPAN4 may be a key regulatory mechanism for the generation of migrasomes.

### 3.8 The changes in cilia–migrasome association are related to breast cancer metastasis and fracture healing

The above results found that EMT is a key signaling pathway of cilia–migrasome association, which has been confirmed to be a key signaling pathway for tumor occurrence and metastasis, tissue regeneration and repair [18, 19]. Given that cilia–migrasome association is abnormal in a variety of tumor tissues, we explored whether cilia–migrasome association is related to tumorigenesis and metastasis in rat DMBA breast cancer model. The primary and metastatic breast cancer tissues of rats were obtained and immunostained for ARL13B and PIGK. As a result, we found that the density and intensity of ARL13B and PIGK positivity in primary tumors were lower than those in metastatic tumors (**Figures 8A, B**), indicating that the decreased level of cilia-migrasome was related to tumor metastasis.

**Figure 8.**
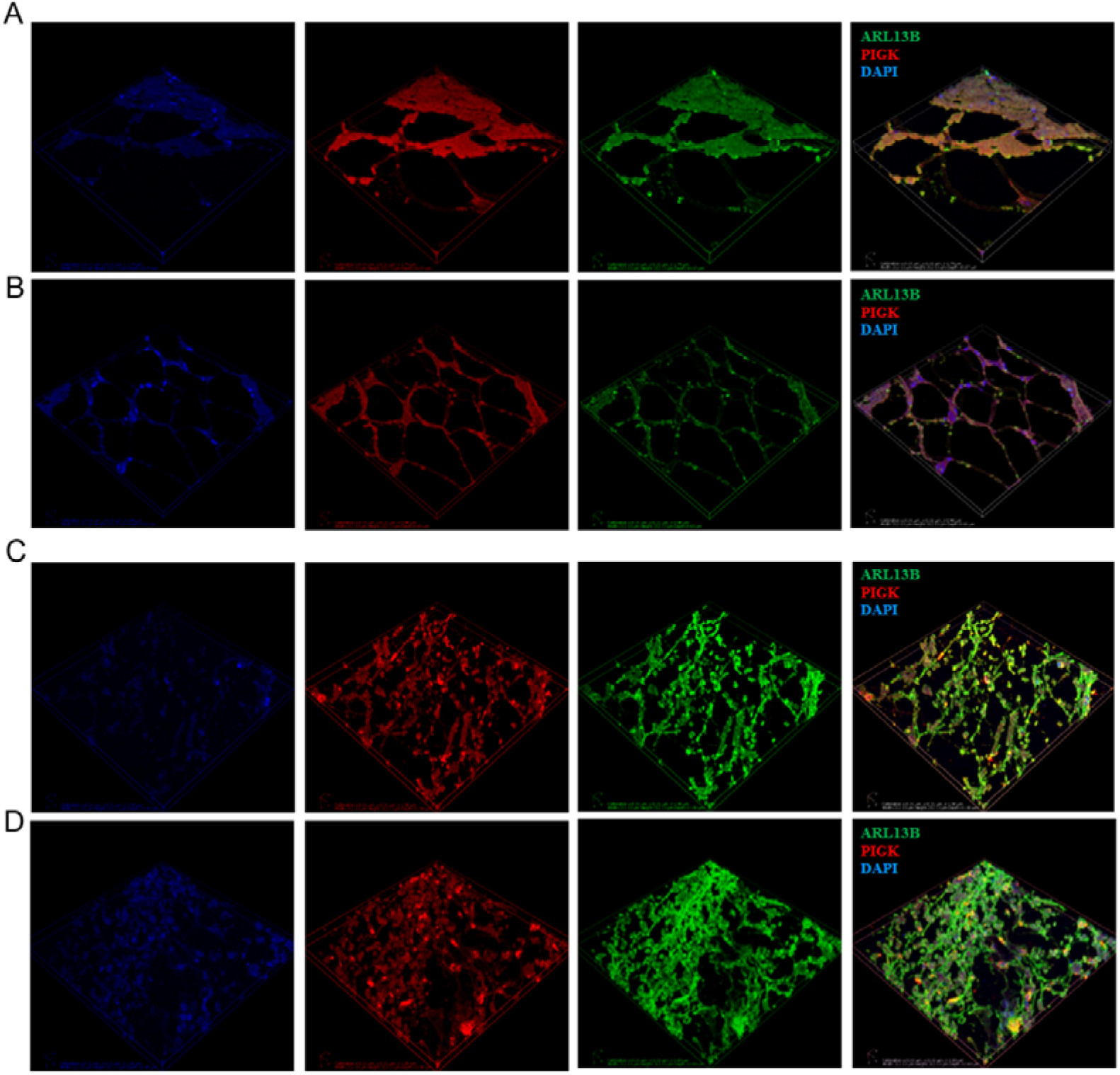
The changes in cilia–migrasome association during breast cancer metastasis and fracture healing. (A,B) Focusing microscope 3D image of immunostaining of ARL13B and PIGK in primary breast cancer (A) and metastatic breast cancer (B). (C, D) Confocal microscopy 3D images of immunostaining ARL13B, of PIGK in 3-day (C) and 5-day (D) mouse fracture repair samples.

Our previous study found that migrasomes are involved in the process of fracture repair. Here we explored whether the change of cilia–migrasome association occurs during fracture healing. By immunostaining ARL13B and PIGK in 0-, 3-, and 5-day mouse fracture samples, we found that ARL13B and PIGK were positively correlated during fracture healing (**Figures 8C, D**), suggesting the potential association of cilia-migrasome in tissue regeneration and repair. The changes in cilia–migrasome association may also be a new perspective to explain tumorigenesis and tissue regeneration.

## 4 Discussion

In this study, we, for the first time, investigated the cilia–migrasome association and found that migrasome are derived from cilia. We further propose that migrasomes may represent a vesicular form of cilia. This interpretation is supported by the following observations: 1.Bioinformatics convergence: migrasome marker expression correlates with cilia-related gene set scores; 2.Morphological evidence: High-resolution SEM reveals cilium-like structures with enlarged tips adjacent to migrasome-like vesicles.In addition, the cilia–migrasome association was found to be related to tumor metastasis and fracture healing. Our data indicates that the cilia–migrasome association not only explains the source of migrasome, but also provides a new perspective to explain the pathological mechanism of human diseases and tissue regeneration and repair.

The ciliary membranes of all cilia contain specific receptor and ion channel proteins that initiate signaling pathways that control motility and/or link mechanical or chemical stimuli, including sonic hedgehog and growth factors, into intracellular transduction cascades that regulate differentiation, migration, and cell growth during development and into adulthood. When cilia fail to form properly or are insufficient in number or cilia are dysfunctional, multiple defects occur, leading to the development of disease. This phenomenon has been found in many common motor system diseases such as osteoarthritis, intervertebral disc herniation, scoliosis, and the occurrence and metastasis of tumors [20]. However, the research and targeted therapy of cilia abnormalities are not satisfactory, because cilia are ubiquitous in most cells of the human body, and our understanding of different physiological and pathological conditions is still lacking [21, 22]. Moreover, because cilia are bound to the cell membrane and have compartmentalized structures specialized for signal transduction, it is difficult to extract cilia and to understand the chemical structure and molecular changes of cilia in different pathological processes. Therefore, cilia-derived extracellular vesicles provide an effective way to understand the role of cilia, which has been studied in model organisms *C. elegans* and *Chlamydomonas* [23, 24].

Shedding of EVs from ciliated cells and EV-cilia incidence relation are evolutionarily conserved phenomena, but the incidence relation between cilia and EVs and the basic biological characteristics of EVs remain unclear [25]. Some ciliary lesions or ciliopathies are associated with abnormal shedding of EVs or abnormal cilia-EV incidence relation [26, 27], which suggests that cilia may be an important organelle for EV donors or EV targets. In our study, we found that migrasome are cilia-derived vesicles, as evidenced by the significant genetic association between migrasome and cilia, the expression of cilia markers in migrasome and the presence of cilia around migrasome, and the expression of proteins in migrasome and cilia. This is an innovative finding, and these results point to migrasome as vesicles that may mediate ciliary function.

Understanding how cilia function in conjunction with other organelles is important to identify the role of cilia in disease progression. Studies have shown that there is a signal axis between mitochondria and cilia. When mitochondrial dysfunction occurs, the structure and homeostasis of cilia are affected, and the disturbance of communication between them has an important impact on the occurrence of nervous system diseases [28], while renal tubular cells with cilium defects are prone to damage due to mitochondrial defects [29]. These studies reveal the possibility of a signaling network between mitochondria and cilia. However, the subsequent mechanism of the perturbation of the signaling between mitochondria and cilia in disease is still unknown [30]. Although as the recently discovered organelles, migrasomes are important pathways for intercellular information transmission, transport of damaged mitochondria, and maintenance of the quality of the cellular mitochondrial pool [3]. Thus, we hypothesize that migrasome are important mediators of the signaling network between cilia and mitochondria in disease and normal physiological processes. However, this speculation remains to be further explored. Nevertheless, our study is the first to reveal the underlying cilia–migrasome association, which will help us to understand the interaction network between subcellular structures in cells.

An interesting finding in this work is that EMT is an important regulatory signaling pathway between cilia and migrasome. EMT and ciliogenesis are two cell biological processes that are essential for the development and maintenance of tissue homeostasis [31]. Activation of cellular EMT under physiological and pathological conditions is associated with significant changes in TGFβ, RTK, WNT, Notch, and HH signaling pathways; notably, most of the signaling pathways associated with EMT are regulated by primary ciliated cell signaling [32, 33]. EMT and cilia have complex mutual regulation, that is, EMT regulates primary ciliogenesis and ciliary signaling, while ciliary signaling mutually regulates EMT signaling activation to promote normal development or disease progression [32, 34]. Based on the functional analysis of cilia-migrasome-associated genes, we found that EMT was an important signaling pathway for the regulation of the relationship, and EMT inhibitor-1 treatment significantly suppressed migrasome production in TSPAN4-GFP labeled RAW264.7 cells. With these results, we suggest that the cilia–migrasome association is associated with EMT. This finding not only provides new insights into the etiology of diseases associated with EMT and ciliary dysfunction but also provides a new perspective on the regulation of cell biological functions by EMT. EMT is a signal pathway that has been confirmed to be a key signal pathway for tumorigenesis and metastasis, tissue regeneration and repair [35, 36]. Therefore, in the rat breast cancer model and fracture repair model, ARL13B (a ciliary marker protein) and PIGK (a migrasome marker protein) staining and correlation analysis were performed on tumor metastasis tissues and fracture healing tissues. The results showed that ARL13B and PIGK levels were decreased in metastatic tumors, and ARL13B and PIGK were positively correlated in fracture-healing tissues. These results indicate that cilia–migrasome association is associated with tumor metastasis and tissue regeneration and repair.

In summary, we, for the first time, proposed the phenomenon of cilia–migrasome association and constructed a migrasome proteomic dataset based on a variety of cell types. The finding of cilia–migrasome association is affected by EMT and participates in tumor metastasis and fracture healing providing a new perspective for disease occurrence and tissue regeneration and repair. The understanding of the incidence relation between migrasomes and cilia provides new ideas for uncovering various physiological and pathological mechanisms and exploring therapeutic strategies for “ciliopathy”.

## 5. Declarations

## Acknowledgements

We appreciate so much to a gentleman who prefers to remain anonymous for providing ideas and technical assistance for this article at the very beginning; thanks to Dr. Zhang Qian and Master Cui Haiyang from Professor Wang Mei’s team of the School of Pharmacy of Xinjiang Medical University for their help with the rat breast cancer samples in this article; thanks to Ms. Wang Ping from Urumqi Jingke Xingye Biotechnology Co., Ltd., and Mr. Xiao Xiong and Mr. Zhou Zhengyang from Wuhan Servicebio Technology Co., Ltd. Thanks to the above institutions for providing reagent consumables and technical support services for this article.

## Author contributions

Conception and design of the research: Siyu Zhang, Yi Yang

Acquisition of data: Siyu Zhang, Abudurexiti Kutibiding, Mengci Wang

Analysis and interpretation of data: Siyu Zhang, Abudurexiti Kutibiding, Mengci Wang

Statistical analysis: Siyu Zhang, Abudurexiti Kutibiding

Obtaining funding: Yi Yang

Drafting the manuscript: Siyu Zhang

Revision of manuscript for important intellectual content: Yi Yang

## Data availability statement

The data that support the findings of this study are available from the corresponding author upon reasonable request.

---

**Supplemental data 1.**
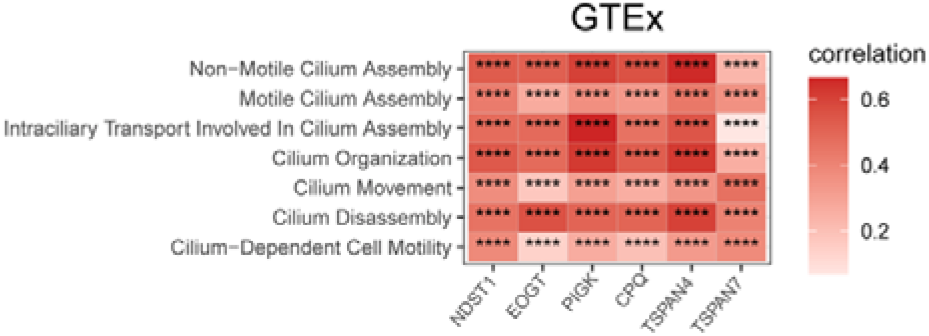
Cilia–migrasome association in various healthy tissues. Spearman correlation analysis was performed between the expression of migrasome-associated markers (“NDST1”, “EOGT”, “PIGK”, “CPQ”, “TSPAN4”, “TSPAN7”) and the ciliary major functional gene set scores, respectively, in total normal tissue samples from the GTEx dataset.

**Supplemental data 2.**
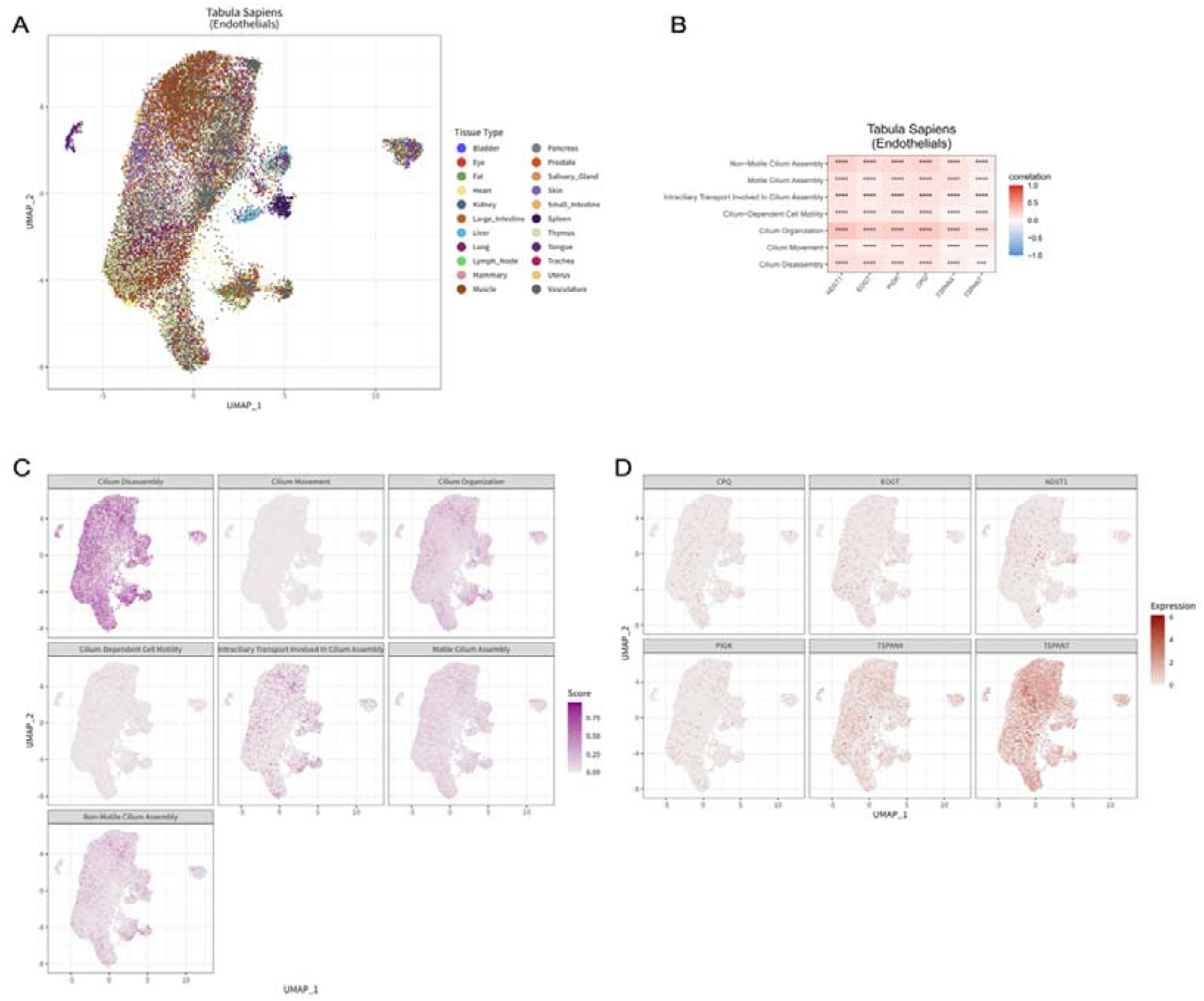
Cilia–migrasome association in endothelial cells. (A) UMAP plot of epithelial cells derived from normal tissues and organs. (B) Spearman correlation analysis was performed between the expression of migrasome-associated markers (“NDST1”, “EOGT”, “PIGK”, “CPQ”, “TSPAN4”, “TSPAN7”) and the ciliary major functional gene set scores, respectively. (C) UMAP plot displaying cilium major functional gene set scores distributing in endothelial cells. (D) UMAP plot displaying migrasome-associated markers in endothelial cells.

**Supplemental data 3.**
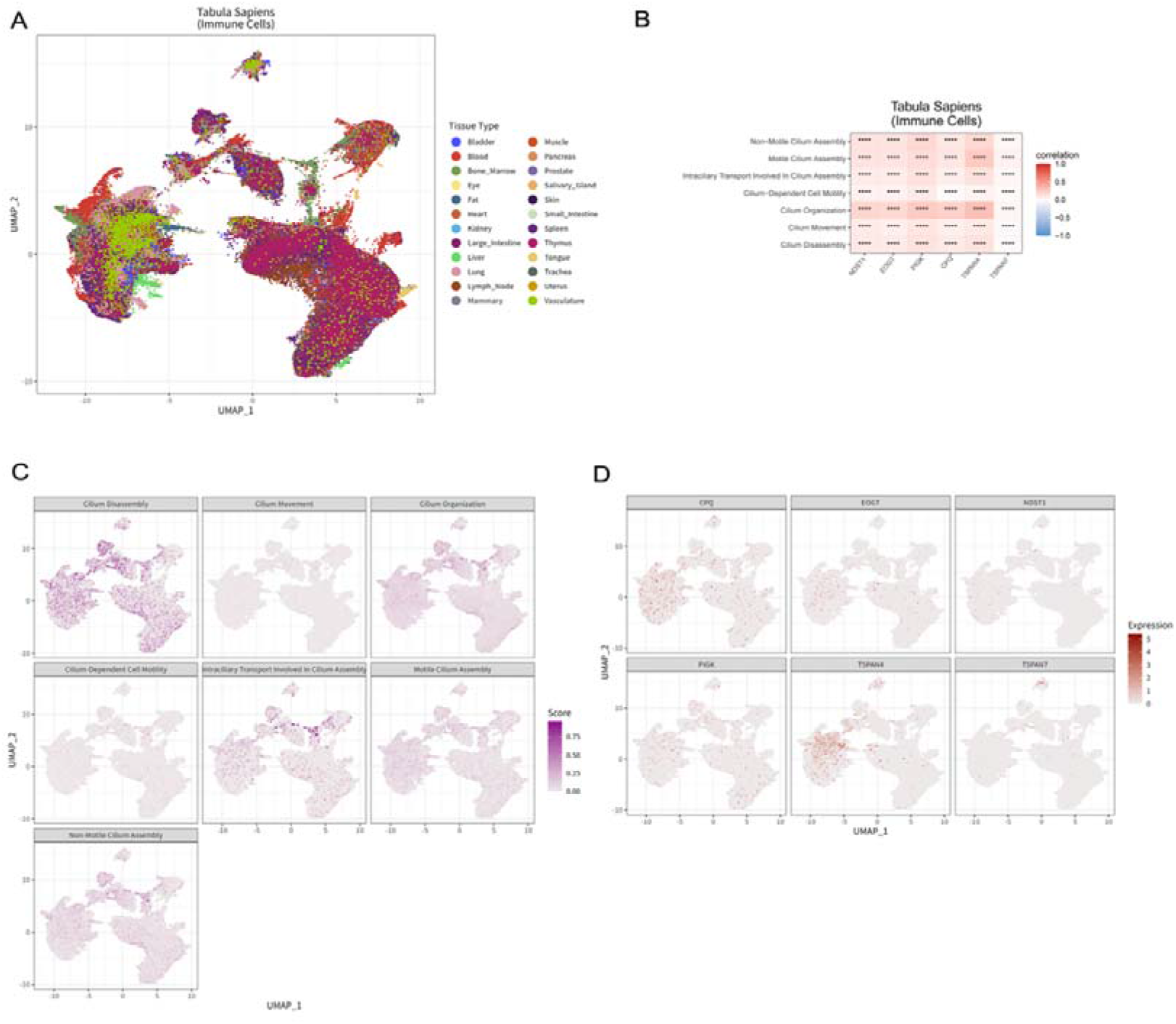
Cilia–migrasome association in immune cells. (A) UMAP plot of epithelial cells derived from normal tissues and organs. (B) Spearman correlation analysis was performed between the expression of migrasome-associated markers (“NDST1”, “EOGT”, “PIGK”, “CPQ”, “TSPAN4”, “TSPAN7”) and the ciliary major functional gene set scores, respectively. (C) UMAP plot displaying cilium major functional gene set scores distributing in immune cells. (D) UMAP plot displaying migrasome-associated markers in immune cells.

**Supplemental data 4.**
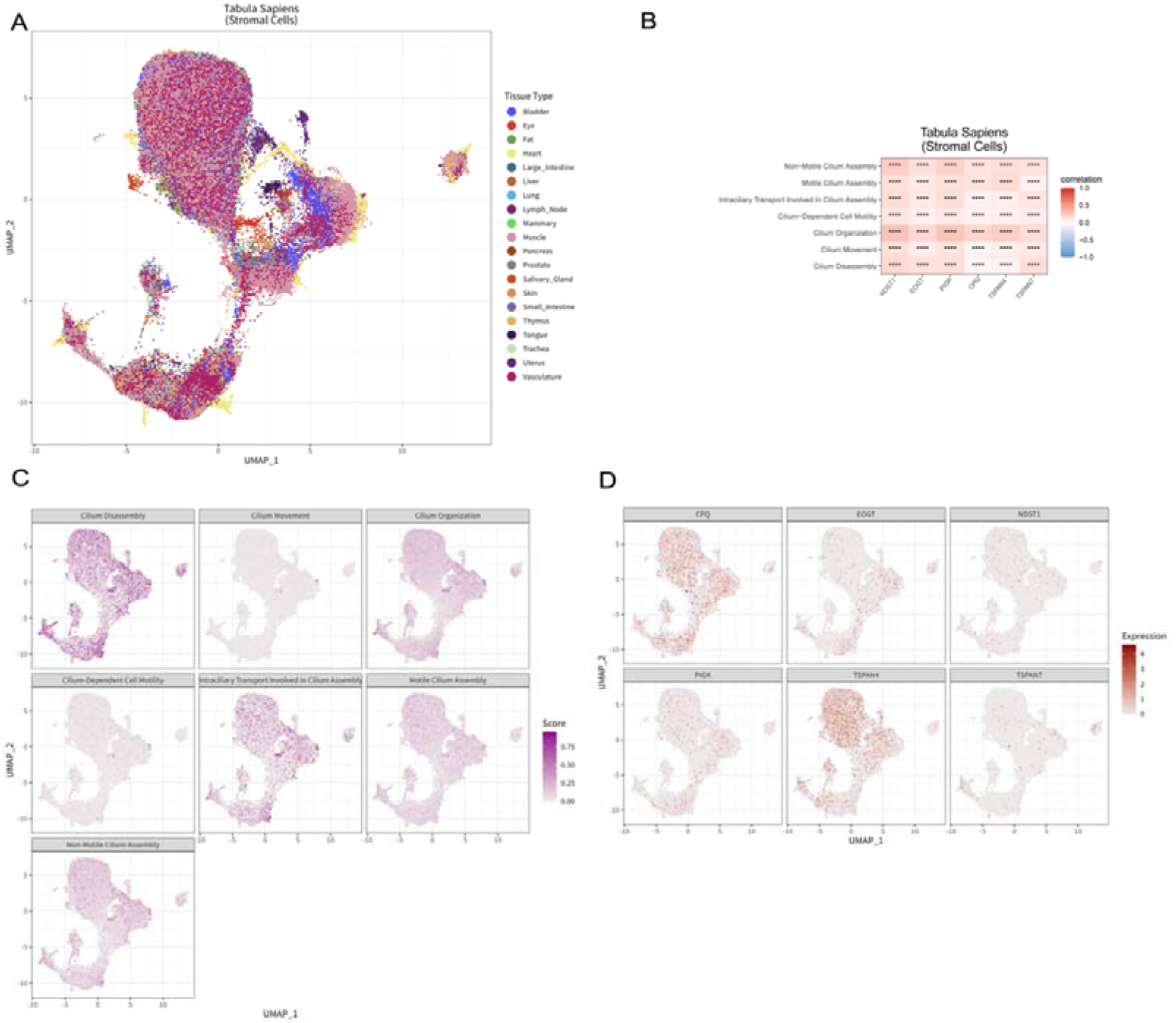
Cilia–migrasome association in stromal cells. (A) UMAP plot of epithelial cells derived from normal tissues and organs. (B) Spearman correlation analysis was performed between the expression of migrasome-associated markers (“NDST1”, “EOGT”, “PIGK”, “CPQ”, “TSPAN4”, “TSPAN7”) and the ciliary major functional gene set scores, respectively. (C) UMAP plot displaying cilium major functional gene set scores distributing in stromal cells. (D) UMAP plot displaying migrasome-associated markers in stromal cells.

**Supplemental data 5.**
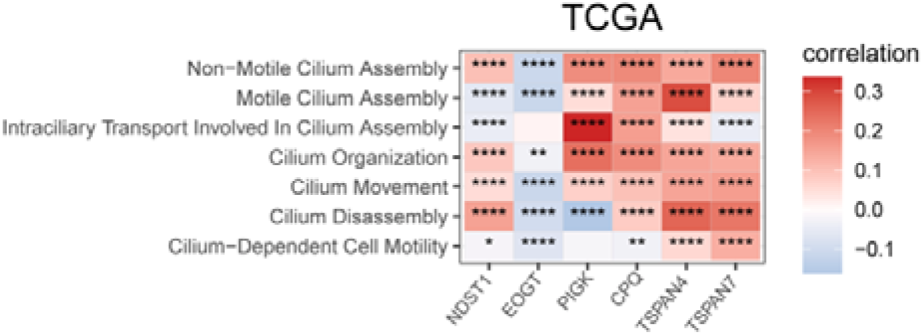
Cilia–migrasome association in various tumor tissues. Spearman correlation analysis was performed between the expression of migrasome-associated markers (“NDST1”, “EOGT”, “PIGK”, “CPQ”, “TSPAN4”, “TSPAN7”) and the ciliary major functional gene set scores, respectively, in total tumor tissue samples from the TCGA dataset.

